# The Cornea Harbors a Tricellular Neuro-Immune Niche that Underpins Touch Sensation

**DOI:** 10.1101/2025.07.28.664937

**Authors:** S Littleton, EM Jacob, Yu Chen, EG O’Koren, NJ Reyes, R Matthew, J Kalnitsky, S Komai, A Nguyen-Thai, S Parikh, W Perez, S Mitra, JL Neff, F Ginhoux, VL Perez, A Matynia, DR Saban

## Abstract

Piezo2 is a mechanosensitive ion channel essential for touch and proprioception, yet the mechanisms that maintain this sensory modality in adult tissues are unknown. Using multiphoton imaging of the cornea in live mice, we discovered that the *Cx3cr1^Cre^* locus targets not only macrophages, but also a distinct subset of nerves. Spatial-RNAseq resolved that *Cx3cr1^Cre^*-driven labeling was uniquely enriched in Piezo2-expressing neurons, a result of temporal *Cx3cr1* expression during development. Through lineage tracing, scRNAseq, and imaging, we identified a novel tripartite cellular niche at the epithelial basement membrane, comprised of monocyte-derived macrophages, nerves, and Schwann cells. Additional scRNAseq and genetic studies revealed that Schwann cell-derived *IL34* maintained corneal macrophages. Through pharmacologic and genetic perturbations, we also demonstrate corneal macrophages selectively maintained the structure-function of Piezo2-enriched nerve endings, with disruption of this niche causing specific deficits in mechanosensation while preserving other sensory modalities. Altogether, we describe a novel tricellular niche in the cornea required for Piezo2-mediated touch sensation, suggesting new directions for investigating mechanosensory circuits including proprioception.

## INTRODUCTION

The proper neuronal sensing of mechanical stimuli is essential for organism survival, yet our understanding of how these sensory circuits are maintained in adult tissues remains incomplete. While neurons were traditionally considered the primary mediators of mechanosensation, evidence now suggests crucial roles for non-neuronal cells beyond glia. For example, in pathological states, bidirectional neuroimmune communication is well established – nerves orchestrate immune responses^1,2,3,4,5,6,7,8^ including macrophage activity, while macrophages influence neuron structure and function^9,10,11^. Less is known in healthy tissue although it has been appreciated that macrophages maintain close physical relationships with certain nerves, and there is compelling evidence supporting a macrophage role in neurophysiology in gut, dermis, and brown adipose tissue^12,13,14,15,16,17^. However, the specific neuronal populations supported remain poorly defined, and nothing is known in this regard for the physiology of Piezo2 mechanosensation.

Piezo proteins are a family of mechanically gated ion channels whose discovery explained how touch is detected and transduced at the molecular level^18^. Piezo2 is expressed in Aδ-fibers (rapidly conduct touch) and some C-fibers (slowly transmit pain, temperature, and itch)^19^. The Piezo2 protein forms a three-part structure whose propeller blades deform the cell membrane, translating mechanical tension into channel opening through an intracellular relay system^20^. In mice, Piezo2 is essential for a variety of mechanosensory responses, including aspects of light touch, and proprioception^21,22^. Likewise, humans with rare mutations in the *PIEZO2* gene have profound mechanosensory deficits that include a loss of the sense of proprioception^23^. Despite these insights, the specific cellular interactions that support Piezo2 nerve function in the steady state remain largely unexplored^24,25,26^.

Here, using two photon imaging of the cornea in live mice, we show that the *Cx3cr1^Cre^* locus, previously used to study macrophages, also marks Piezo2-enriched nerves, with these macrophages and nerves existing in contact throughout the tissue. We show that monocyte-derived corneal macrophages form a tri-cellular niche with nerves and Schwann cells at the epithelial basement membrane. In this unique niche, IL-34 is expressed by Schwann cells and maintains macrophages, while macrophages maintain the structure-function of Piezo2-enriched mechanosensory nerve endings. Altogether, these findings reveal a novel immune cell contribution to sensory function under physiologic conditions and open new avenues for exploring the intricate cellular interactions that underpin sensory function.

## RESULTS

### Monocyte-derived macrophages populate the adult cornea and partake in a site-specific arrangement with nerves and Schwann cells

All tissues are seeded during development by macrophages, derived from the yolk sac (YS) or fetal liver (FL), but many are eventually repopulated through bone marrow-derived definitive hematopoiesis in adults^27^ (with exception of the brain and other specific tissues). Here, we studied corneal macrophage ontogeny using three separate lineage tracers. First, we assessed the YS primitive macrophage fate mapper *Runx1^Mer-Cre-Mer^; Rosa^R26R-eYFP^* embryos by pulsing with 4-hydroxytamoxifen (4-OHT) at embryonic day 7.5 (E7.5)^28^. Whole mount corneas from progeny mice at 8 weeks were analyzed by confocal microscopy for the macrophage marker Iba-1. As a positive control, we analyzed the retina for microglia; as a negative control, we analyzed cornea of age matched *R26^LSL-fGFP^* mice ^29^. We found little to no YFP+ macrophages in the cornea, indicating that adult corneal macrophages are not YS-derived (Sup Fig 1A-B). Likewise, historical data allowed us to also rule out long-term presence of FL-derived macrophages based on our previous work demonstrating that cornea of *Cx3cr1^CreER^; R26^LSL-fGFP^* mice are absent of fGFP macrophages in the cornea one-year post-tamoxifen pulse^29^. Focusing on definitive hematopoiesis, we took advantage of *Ms4a3^Cre^; R26^LSL-tdTom^* mice, which trace derivatives of granulocyte monocyte progenitors (GMP)^30^. Separately, we also analyzed *Flt3^Cre^; R26^LSL-tdTom^* mice, which labels adult hematopoiesis-derived cells^31^. In both mouse strains, confocal analysis of whole mount cornea and immunolabeling against the macrophage marker Cd206 revealed colocalization of tdTom+ Cd206+ macrophages throughout the cornea, including central and peripheral regions (Figure 1 A-F). We conclude that adult corneal macrophages are primarily monocyte-derived, consistent with previous reports^29,32,33^.

**Figure 1:**
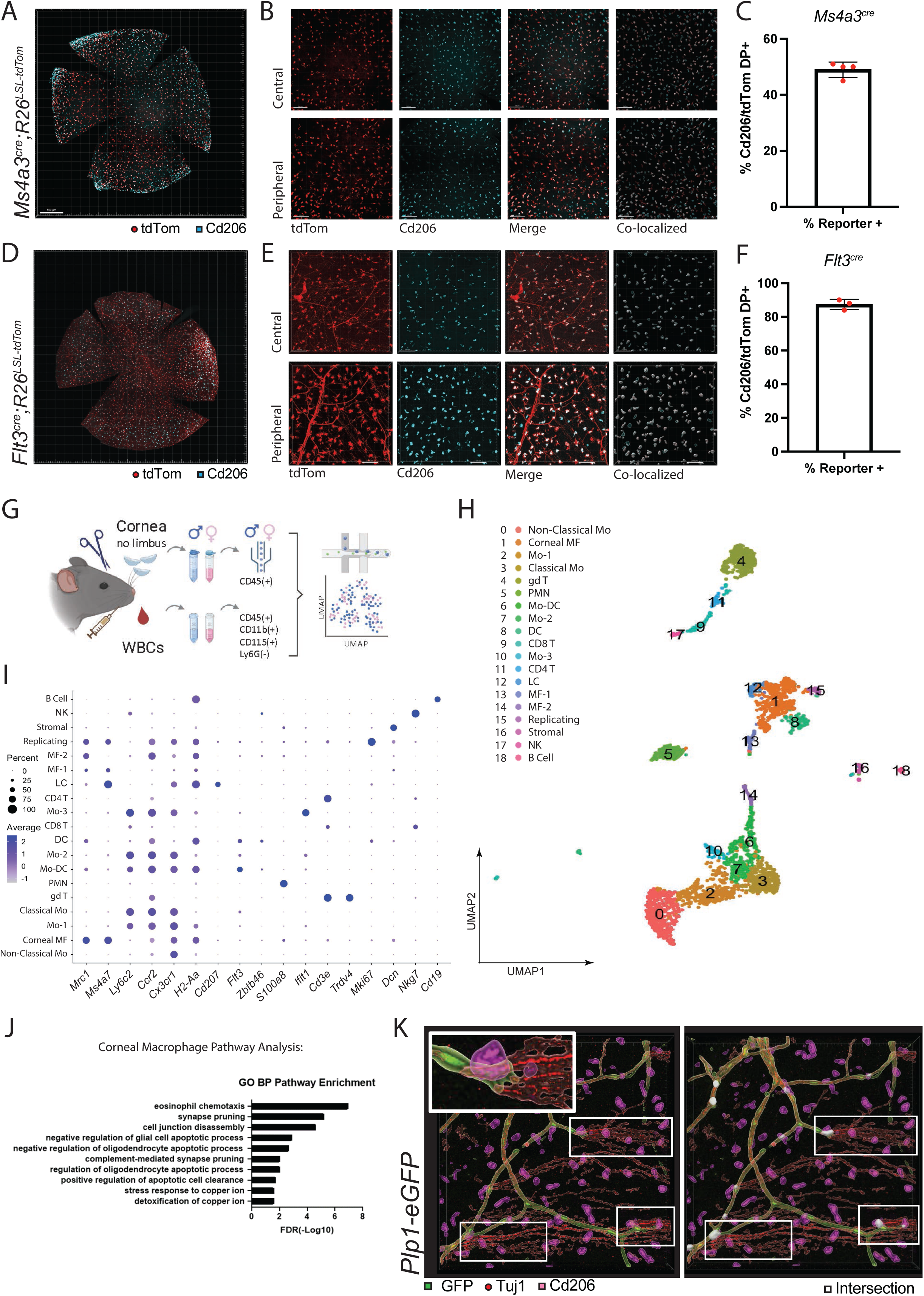
Corneal Macrophages form Tripartite Interactions with Glia and Nerves at Stromal-Epithelial Nerve Penetration Sites. (A) Whole mount confocal microscopy of corneas from 12-week-old *Ms4a3^Cre^; R26^LSL-tdTom^* reporter mice. (B) Representative images from central and peripheral regions, showing tdTom+ monocyte lineage cells (red), Cd206+ macrophages (cyan), merged signals, and rendering of colocalized signals (white). (C) Quantification of Cd206+, tdTom+ double positive cells via confocal microscopy (n=4). (D) Whole mount confocal microscopy of corneas from 9-month-old *Flt3^Cre^; R26^LSL-tdTom^*reporter mice. (E) Representative images from central and peripheral regions, showing tdTom+ cells (red), CD206+ macrophages (cyan), merged signals, and rendering of colocalized signals (white). (F) Quantification of Cd206+, tdTom+ cells via confocal microscopy (n=3). (G) Overview of corneal scRNA-seq methodology (n=98 corneas), including FACS of circulating monocytes from blood. (H) UMAP dimensionality reduction and clustering of scRNA-seq data from corneal immune cells. (I) Dot plots of enriched marker genes for each cell cluster in the cornea. (J) Bar graphs of top-ranked pathways (Biological Process) from Gene Ontology (GO) pathway enrichment analysis. (K) *Ex-vivo* corneal confocal microscopy of *Plp1-eGFP* Schwann cell reporter mice showing rendered intersection (white) of macrophage (purple), Schwann cell (green), and nerve (red) volumes at stromal-epithelial nerve penetration sites (representative of at least n=3 independent experiments). A/D scale bar = 200 µm; B/E scale bar = 100 µm.

To profile corneal macrophages, we next performed single-cell RNA-seq (scRNA-seq) using the 10× Genomics platform. FACS was used to isolate live Cd45+ cells from corneal single cell suspensions (n=25 male; n=25 female mice). Special care was made to exclude the limbus, thus avoiding contamination from circulating leukocytes in the vascular arcade and adjacent conjunctival tissues. Separately, we sorted Cd45+ Cd115+ Cd11b+ Ly6g-cells from circulating blood, which, by definition, contain monocytes that give rise to corneal macrophages (Figure 1G). Cells were pooled by sex for library preparation. A total of n=2800 cells passed quality control (QC), and unbiased hierarchical clustering yielded 19 clusters broadly characterized as lymphoid, corneal myeloid, and circulating monocytes (Figure 1H-I). No major differences could be observed across sex (Supplemental Figure 1C). Lymphoid clusters were characterized by gamma-delta T cells, which are sparse in the cornea^34,35^, with small contributions from CD4 and CD8 T cells, as well as innate lymphoid cells (ILCs). Classical and non-classical monocytes were characterized by their expression of *Ccr2*, *Ly6c2* and *Cx3cr1*, *Nr4a1,* respectively (Supplemental Figure 1D). Macrophages were identified by their high expression of *Ms4a7* and *Mrc1* (Supplemental Figure 1E). Langerhans cells and dendritic cells were identified due to their high expression of *Cd207* and *Zbtb46*, respectively (Figure 1H-I). *Mki67* and *Top2a* suggested the possible presence of a replicating macrophage population within mouse cornea (Figure 1H-I).

Our next aim was to interrogate the functional significance of corneal macrophages on the basis of their transcriptomes. To predict the terminally differentiated macrophage cluster for which to examine, we performed trajectory analysis using both Monocle3 and Velocity via scVelo^36,37,38^. Our results showed that both packages predicted that the monocyte-derived cell adaptation to the corneal niche terminates at cluster 1, corneal macrophage (Supplemental Figure 1E). Next, we performed Gene Ontology (GO) Enrichment Analysis^39^. Our results indicated that five of the top ten pathways predicted neuro-immune interactions. Two such pathways involved interactions with neurons, including *synapse pruning* and *complement-mediated synapse pruning* (Figure 1J). The other three pathways involved interactions with glia, including *negative regulation of glial cell apoptotic process*, *negative regulation of oligodendrocyte cell apoptotic process*, and *regulation of oligodendrocyte cell apoptotic process* (Figure 1J). These results suggest that corneal macrophages play a significant role in the maintenance of glia and neurons.

To validate our GO results, we performed immunofluorescence confocal microscopy of mouse corneas. While the cornea does not contain oligodendrocytes, it is replete with myelinating and non-myelinating Schwann cells^40,41^. As such, we took advantage of *Plp1-eGFP* reporter mice that label Schwann cells^42^ (<u>Supplemental Video 1</u>). In addition, we immunolabeled nerves with beta-3 tubulin (Tuj1) and macrophages with Cd206. We used Imaris Surface models for co-localization analysis, enabling us to identify events in which GFP (Schwann), Tuj1, and Cd206 cell volumes intersected. The only locale in which all 3 cell types intersected was at the epithelial basement membrane (Figure 1K, <u>Supplemental Video 2</u>). This location represents penetration points where nerves, which project from the ophthalmic branch (V1) of the trigeminal ganglion, transition from the corneal stroma to the epithelium. At this transition, the nerves shed their glial cell ensheathment^43,44,45,46,47,48,49,50^. Altogether, our genetic, sequencing and spatial analyses identified a unique tripartite cellular niche comprised of nerves, glia, and macrophages at the epithelial basement membrane.

### IL-34 is expressed by Schwann cells and contributes to corneal macrophage maintenance

Our gene enrichment analysis predicted a role for corneal macrophages in the maintenance of glia. As such, we asked whether macrophage depletion would cause detectable Schwann cell defects. Our approach was to longitudinally follow corneal Schwann cell morphology after macrophage depletion. We used anti-CSF1R antibody mediated depletion of macrophages in *PLP1-eGFP* mice (Figure 2A). We analyzed the cornea longitudinally via intravital two-photon (2P) microscopy (Figure 2B-C). Confocal microscopy was used post-mortem to confirm macrophage depletion (Figure 2D). Images captured by 2P from live mice were analyzed by Imaris software using Surfaces model to render central corneal Schwann cell volumes. Results showed that there were no significant differences between baseline, depleted, and control groups (Figure 2E). Likewise, ex-vivo analyses further validated a gross lack of changes in PLP1+ Schwann Cells (Figure 2D). Taken together, our results suggest macrophages are not critical for the maintenance of Schwann cell morphology.

**Figure 2:**
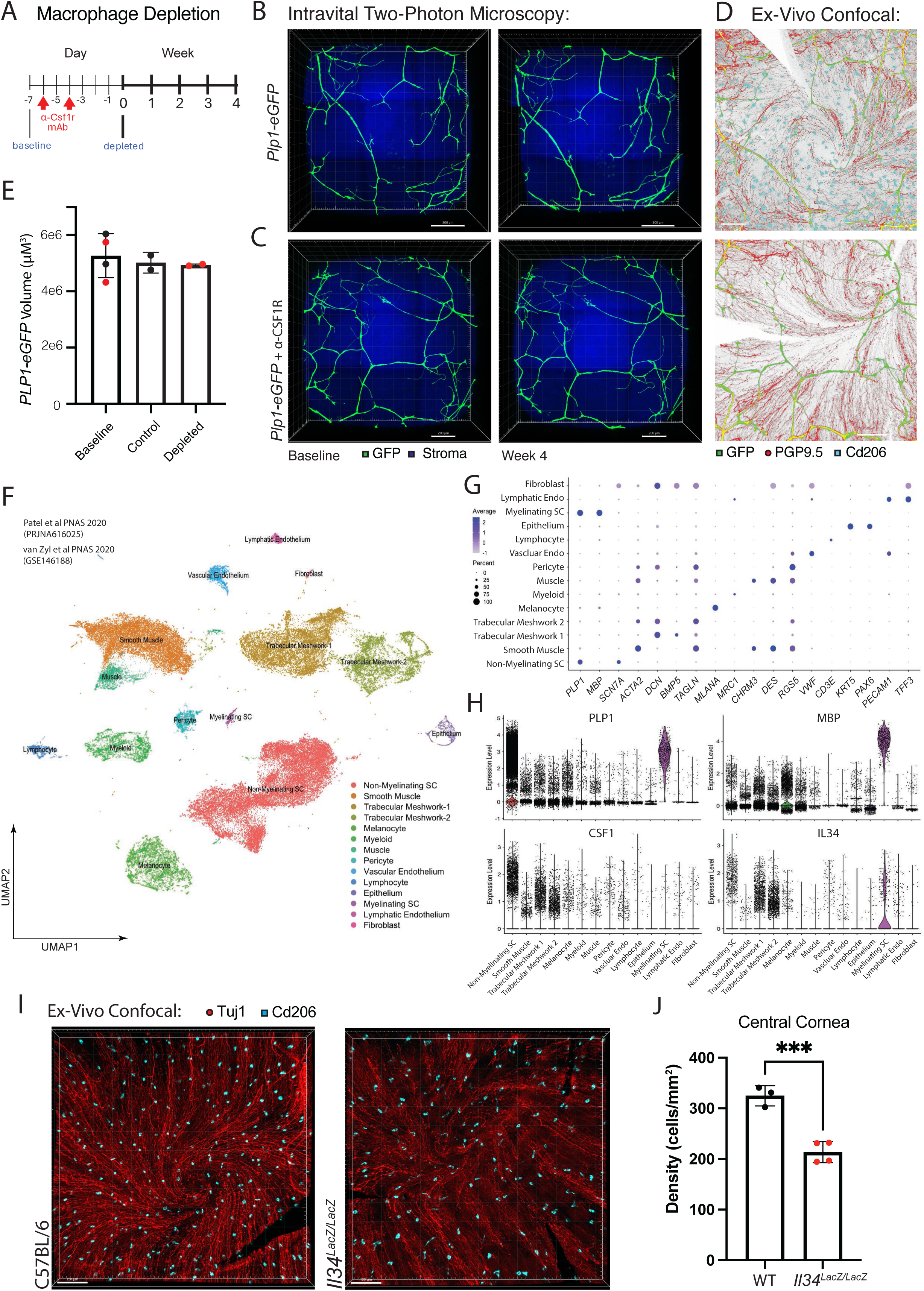
The Functional Significance of Schwann Cell-Macrophage Interactions. (A) Schematic of macrophage depletion experiment. (B) Intravital multiphoton imaging of *Plp1-eGFP* corneas, with second harmonic generation (SHG) of the corneal stroma (blue). (C) Intravital images of macrophage-depleted Plp1-eGFP mice. (D) Ex-vivo confocal imaging of control or macrophage depleted, whole mount *Plp1-eGFP* corneas, showing Cd206+ macrophages (purple) and Tuj1+ nerves (red). Identical field of view detected between imaging modalities. (E) Quantification of Plp1+ Schwann cell volume in live mice, at baseline and following macrophage depletion (n=2 control, n=2 anti-CSF1R). (F) Integration and re-clustering of publicly available human anterior segment single-cell RNA sequencing data. (G) Dot plots depicting enriched marker genes for each cell cluster from the integrated human anterior segment dataset. (H) Violin plots of *CSF1* and *IL34* expressed in both myelinating and non-myelinating Schwann cell populations. (I) Whole mount confocal images of WT and *Il34*-deficient mouse corneas, showing Tuj1+ nerves (red) and Cd206+ macrophages (cyan). (J) Quantification of corneal macrophage numbers in *Il34*-deficient mice (n=3 WT, n=4 KO). Data shown as mean ± SEM, analyzed using one-way analysis of variance (ANOVA) for repeated measures of Schwann cell volume or t-tests for quantifying macrophage numbers. ∗p < 0.05; ∗∗p < 0.01; ∗∗∗p < 0.001; ∗∗∗∗p < 0.0001; ns, not significant. B scale bar = 300 µm; C/D scale bar = 200 µm; I scale bar = 150 µm.

We next tested the converse scenario—whether Schwann cells contribute to corneal macrophage maintenance. Our approach was to reanalyze publicly available datasets from human anterior segment tissues^51,52^ (Figure 2F). Using marker genes, we were able to assign cell types (Figure 2G), with *PLP1*, *MBP,* and *SNC7A* identifying myelinating and non-myelinating Schwann cell clusters (Figure 2H). Interestingly, we found unique enrichment of macrophage trophic factors *CSF1* and *IL34* within Schwann cell clusters (Figure 2H). We note trabecular meshwork clusters also exhibited high expression of these genes, but these cells are not part of the cornea. This suggests a possible role for Schwann cells in corneal macrophage maintenance.

Based on these results, we then asked how macrophages are maintained locally in the cornea. We addressed this question using knockin *Il34^Lacz/LacZ^*mice, which are *Il34* deficient^53^. Whole mount corneas from wildtype versus *Il34^Lacz/LacZ^* mice were immunolabeled for Cd206 and Tuj1 and Cd206+ macrophages were quantified in the context of Tuj1 corneal nerves (Figure 2I). Our results revealed that *Il34^Lacz/LacZ^*mice had significantly less Cd206+ macrophages in the central cornea, suggesting IL-34 is important for their maintenance (Figure 2J). Taken together, these results suggest that Schwann cells play a key role in the local maintenance of corneal macrophages via IL-34, whereas macrophages do not appear to contribute to the maintenance of Schwann cell morphology.

### Corneal macrophages maintain a unique population of corneal nerves

We next asked whether macrophages contribute to nerve maintenance in the normal adult cornea. To address this question, we took advantage of a reporter system that labels both nerves and macrophages simultaneously – thus enabling longitudinal intravital 2P imaging of the cornea. We identified that *Cx3cr1^Cre^* mice drive this dual reporter feature, as either BAC transduced *Cx3cr1^Cre^; R26^LSL-tdTom^* mice^54,55,56^ (<u>Supplemental Video 3</u>) or knockin *Cx3cr1^Cre/+^*; *R26^LS-fGFP^* mice^57^ (Sup Fig 2A). We note that corneal nerve labeling was absent in adult *Cx3cr1^eGFP/+^* mice^58^ and adult tamoxifen pulsed *Cx3cr1^CreERT^*^2^; *R26^LSL-tdTom^* mice^57^ (Sup Fig 2B). These results indicate that *Cx3cr1* is not expressed in adult corneal nerves, therefore suggesting that *Cx3cr1^Cre^* nerve labeling is likely due to a temporal expression during development.

Using *Cx3cr1^Cre^; R26^LSL-tdTom^* adult mice, we depleted macrophages with anti-CSF1R mAb and followed the cornea by longitudinal intravital 2P imaging. Within 96 hours of complete macrophage depletion, we found a distinct loss of *Cx3cr1^Cre^*-labeled nerve leashes impacting the whorl-like patterning (Figure 3A) (Supplemental Video <u>4</u> & <u>5</u>). This atrophy was observed within the subbasal plexus – located between the epithelial basement membrane and the basal epithelium (Figure 3B). The significance of this location is that it is directly upstream of nerve penetration points, exactly where we found the tripartite interacting macrophages. Given that macrophage depletion results in the atrophy of *Cx3cr1^Cre^*-labeled nerve leashes, we can conclude that macrophages support corneal nerves, potentially at the level of the subbasal plexus.

**Figure 3:**
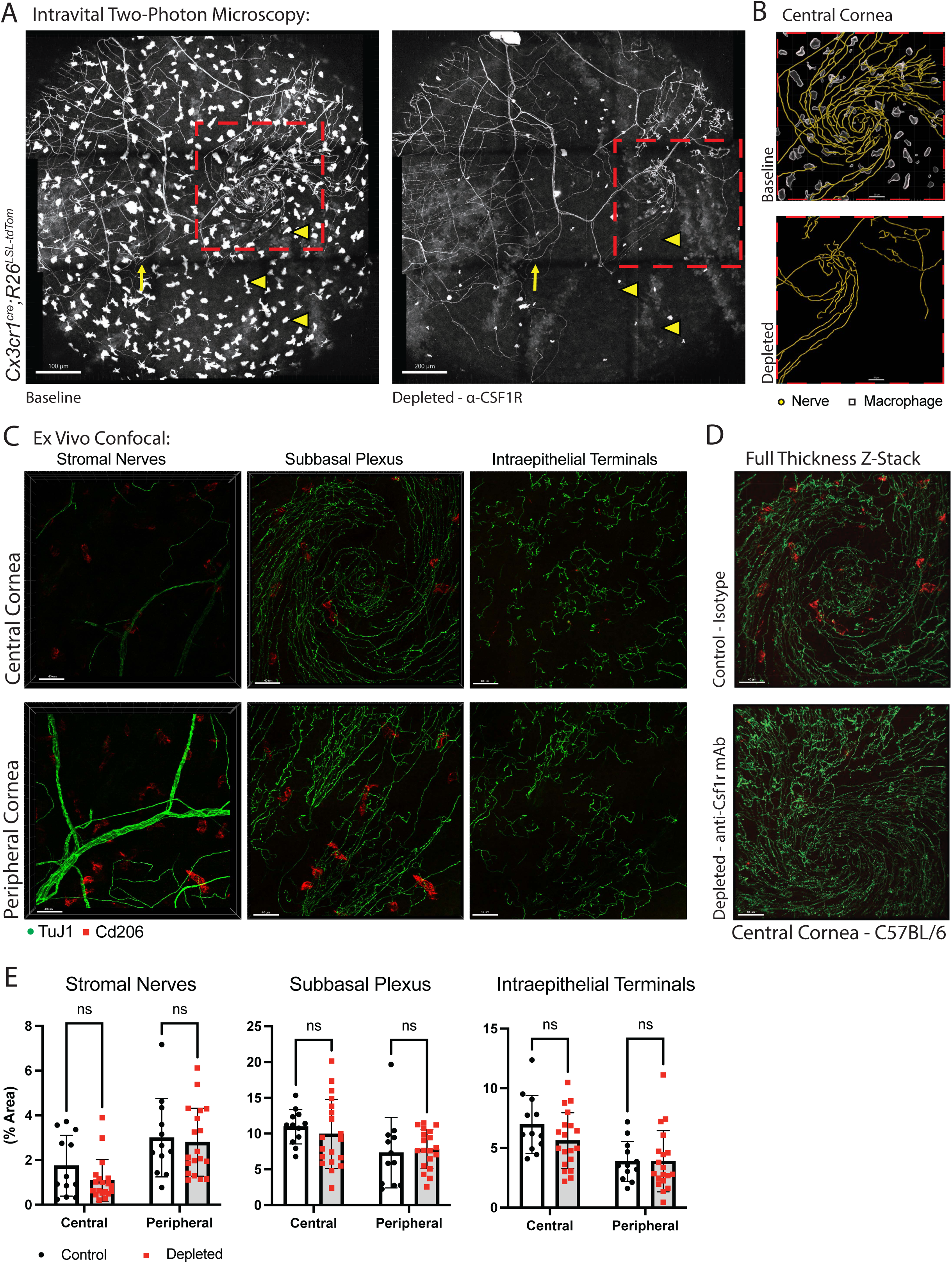
Macrophages Maintain a Select Population of Corneal Nerves. (**A**) *Cx3cr1^cre^;R26^LSL-tdTom^* visualized by intravital multiphoton microscopy. Nine-tile z-stacks were acquired and stitched to visualize corneal nerves and immune cells (yellow arrowheads depict depleted macrophages; yellow arrow marks a degenerating nerve leash; red box depicts an identical field of view between baseline and depleted; scale bar = 100 µm). (B) Imaris renderings of baseline and macrophage-depleted central corneas (see supplemental videos for dynamic imaging). Scale bar = 50 µm. (C) Representative ex-vivo confocal images of central and peripheral stromal, sub-basal plexus, and epithelial regions, showing nerve (green) and macrophage density (red). Scale bar = 40 µm. (D) Representative whole mount images of the central cornea from isotype and anti-CSF1R treated mice. Scale bar = 40 µm. (E) Quantification of corneal nerve density from central and peripheral regions of stromal, sub-basal plexus, and epithelium (n=12 control; n=19 depleted). Data shown as mean ± SEM, analyzed using t-tests. ns, not significant. Data are representative from at least two independent experiments.

We next sought to better characterize the extent of this atrophy, as *Cx3cr1^Cre^*only labels a small fraction of corneal nerves. Our approach was to harvest macrophage-depleted corneas for Tuj1 immunolabeling and assess nerves via confocal microscopy within the stroma, subbasal plexus, and epithelial termini of the central and peripheral cornea (Fig 3C-D). Surprisingly, we found no significant changes in the depleted cornea by assessing % area of all Tuj1+ nerves, although we could detect a trend in the central cornea (Figure 3E). To further assess this observation, we turned back to longitudinal intravital 2P imaging, this time using CGRP-GFP mice^54,59^, as most corneal nerves express CGRP. Again, minimal to no changes were observed in the nerve patterning following macrophage depletion (Sup Fig 2C). As we were only able to detect significant remodeling in *Cx3cr1^Cre^*-labeled corneal nerves, we conclude that macrophages support this unique population, and the support occurs at the subbasal plexus.

### *Cx3cr1^Cre^*-labeled somata in the trigeminal ganglia are *Piezo2-*enriched

To determine the identity of *Cx3cr1^Cre^*-labeled corneal nerves, we employed GeoMx whole transcriptome atlasing, spatial sequencing platform on trigeminal ganglia of *Cx3cr1^Cre^; R26^LSL-tdTom^* mice. Trigeminal ganglia (TG) were dissected and inked to assist in proper orientation during embedding (Figure 4A-B). We placed 1 section per mouse, fitting 6 sections at equal depth on the capture slide, enabling us to sequence from n=6 mice simultaneously. To confirm our sectioning approach, H&E staining was employed to clearly reveal the nerve somas and axon fibers, allowing the distinction of each of the V-regions (Figure 4C). Sections were then nuclear stained and Tuj1 immunolabeled to enable proper segmentation for sequencing. To validate our segmentation approach, we drew regions of interest (ROI) within the ganglia and in the axon regions (Sup Fig 3A-B). Once scanned with GeoMx Digital Spatial Profiler (DSP), sequencing was completed and standardized QC was carried out, yielding n=39 segments from ganglia and n=24 segments from axonal regions. We then validated our segmentation approach by performing hierarchical clustering of each segment (Sup Fig 3C) and dimensionality reduction using PCA (Sup Fig 3D). Results showed that axon segments clustered differently than the soma segments, indicating, as expected, that transcripts expressed are significantly different in somata versus the axons. Accordingly, *Ncmap* (Schwann cell marker) was enriched in the axon ROIs, whereas *Fabp7* (satellite glial cell marker) was enriched in the somata ROIs (Sup Fig 3E).

**Figure 4:**
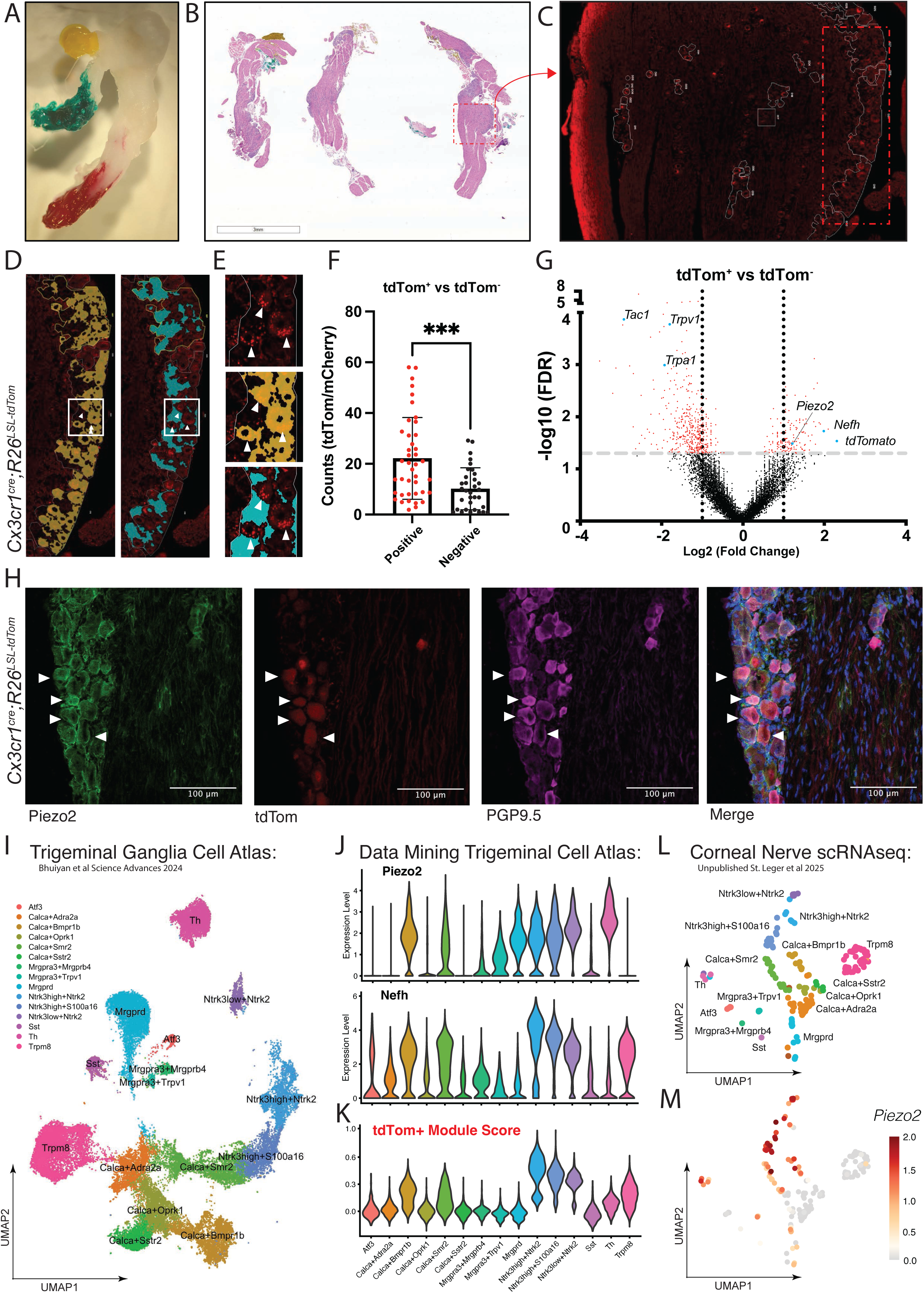
*Cx3cr1^Cre^*Locus Selectively Targets *Piezo2*+ TG Neurons. (A) Dissection of mouse trigeminal ganglia and tissue inking of the V1 (red), V2 (green), and V3 (yellow) branches of the mouse TG nerve to demarcate regions for GeoMx spatial RNAseq analysis. (B) H&E stain of sectioned TG (n=6) used for spatial RNAseq (Scale bar = 3 mm). (C) Representative Region of interest (ROI) selection of V1 region based on tdTom signal from *Cx3cr1^Cre^;R26^LSL-tdTom^*. (D) Region of interest (ROI) segmentation in the V1 region, with tdTom+ signal shown in yellow overlay and tdTom-signal shown in cyan (arrows indicate tdTom+ soma regions). (E) Zoomed-in view of ROI segmentation from panel E. (F) RNAseq reads from tdTom+ and tdTom-segmented regions, demonstrating successful enrichment of sequences from tdTom+ neurons (n=41 tdTom+, n=33 tdTom-). (G) Volcano plot showing DEGs from n=5 tdTom+ and n=5 tdTom-segmented regions. (H) Confocal analysis of TG of *Cx3cr1^Cre^;R26^LSL-tdTom^* mice shows co-labeling of Piezo2, tdTom, PGP9.5 (Scale bar = 100 µm). (I) UMAP of scRNA-seq data from the Harmonized Trigeminal Ganglia Cell Atlas, Bhuiyan *et al.* (J) Violin plots depicting DEGs identified from our spatial-RNAseq of tdTom positive and negative segmented ROIs. (K) tdTom+ module score calculated using genes with positive fold change and FDR > 1.2 projected onto Bhuiyan *et al.* dataset. (L) UMAP of scRNA-seq dataset from corneal retrograde labeled TG somata (unpublished St. Leger *et al.*). (M) Feature plot depicts *Piezo2* expression. Data shown as mean ± SD, analyzed using Welch’s t-test. DEGs were calculated using Multivariate Linear Regression or t-tests using the GeoMx DSP.

We next set out to sequence tdTom+ nerves in the TG of *Cx3cr1^Cre^; R26^LSL-tdTom^* mice. Using the same approach as described above, we were able to clearly detect tdTom+ somata in the TG (Figure 4D), enabling the segmentation of tdTom positive versus negative somata (Figure 4E). Validating our approach, differential gene expression analysis demonstrated significantly enriched *mCherry/tdTomato* expression (Figure 4F). To further optimize our analysis, we focused on well segmented events defined by clear tdTom positive vs negative protein visualization and corresponding *tdTom* gene expression. This approach led to our identification of *Piezo2* enrichment within the tdTom+ population (Figure 4G). We validated these results by immunolabeling for Piezo2 and PGP9.5 expression in TG sections from *Cx3cr1^Cre^; R26^LSL-tdTom^*mice. We were able to show tdTom+ colocalization with Piezo2 and PGP9.5 (Figure 4H).

Given that *Piezo2* encodes a critical ion channel involved in mechanosensation, our results suggested that *Cx3cr1^Cre^*-labeled corneal nerves represent a functional class of sensory neuron^60,61,62^. Going back to our spatial RNAseq results, tdTom+ gene enrichment of *Nefh* may suggest an A-as opposed to C-fiber designation^19^. Consistent with this interpretation, we found *Tac1*, which encodes Substance P and is often co-expressed with *Calca* (CGRP), was strongly downregulated in tdTom+ nerves. To further address this question, we took advantage of the Bhuiyan *et al.* dataset, a cross species harmonized TG cell atlas (Figure 4I). We mined their results for *Piezo2* and *Nefh* expression, as well as the module DEG score of tdTom+ neurons from our spatial RNAseq dataset (Figure 4J-K). Likewise, we performed a separate analysis of an unpublished dataset from cornea retrograde labeled TG somata (St. Leger et al. University of Pittsburg) (Figure 4L-M). In both datasets, we found that *Piezo2*, *Nefh*, and the tdTom module score enrichments all converged on *Nrtk*-expressing populations (Figure 4J-M). According to Bhuiyan *et al.* annotations, which was established using at least one known function ascribed from rodent studies where direct genetic access to the cell type was obtained using knock-in mice expressing Cre recombinase, the enrichments in *Nrtk*-expressing populations we found align with an A-fiber designation^19^. We therefore conclude that *Cx3cr1^Cre^*-labeled somata in the TG represent *Piezo2* expressing A-fiber neurons.

### Corneal macrophages maintain Piezo2 mechanosensory nerves

Our final series of experiments sought to identify the population of nerves supported by corneal macrophages. We took advantage of our findings that these neurons are *Cx3cr1^Cre^* labeled and that *Cx3cr1^Cre^*-labeled somata in the TG are Piezo2 enriched. We performed immunolabeling using anti-Piezo2 antibodies in whole mount cornea. However, despite attempts to optimize fixation and staining strategies, we failed to clearly visualize this population. Next, we turned to *Piezo2*-IRES-CRE-GFP reporter mice^63^, but GFP visualization was too dim to resolve nerve labeling. We then crossed these mice with *R26^LSL-tdTom^* mice but found significant expression of tdTom in non-neuronal cells.

Because we could not identify Piezo2 through our initial approaches, we utilized fluorescent styryl dye (FM1-43), which can enter mechanosensitive channels when they are active, to functionally label Piezo2+ neurons based on their mechanosensitivity^26^. We took advantage of this property to examine the identity and function of the *Cx3cr1^Cre^* labeled corneal nerves. We first established our ability to visualize FM1-43 labeled nerves in the cornea and the range of excitation and emission under intravital 2P microscopy (<u>Supplemental Video 6</u>). Next, we determined whether FM1-43 colocalizes with tdTom in *Cx3cr1^Cre^; R26^LSL-tdTom^* mice. Given that FM1-43 and tdTom have remarkably similar emission properties, we first characterized tdTom+ nerves in corneas prior to FM1-43 administration (Figure 5A), with special attention to also gather signal from lower wavelengths outside of the tdTom spectrum (Figure 5B). Next, FM1-43 was administered i.p. to *Cx3cr1^Cre^; R26^LSL-tdTom^*mice and 24 hours later images of fluorescent signal generated by labeled nerves in the previously negative wavelengths were captured (Figure 5C). Using Imaris, co-labeled nerves were identified by comparing baseline tdTom signals and FM1-43 labeling in the green spectra. With this strategy, we were able to show that many of the tdTom+ nerves were co-labeled with FM1-43 (Figure 5D). Notably, the ability to detect FM1-43 colocalization is limited in these conditions, as the green spectra collected excludes peak emission of the styryl dye, thereby underestimating full labeling. Despite this limitation, our results suggest that tdTom+ FM1-43+ are Piezo2 nerves and exhibit mechanosensitive activity in-vivo. Moreover, we were able to show that FM1-43 labeled somata are Piezo2+ PGP9.5+ in the TG, further validating our findings (Figure 5E).

**Figure 5:**
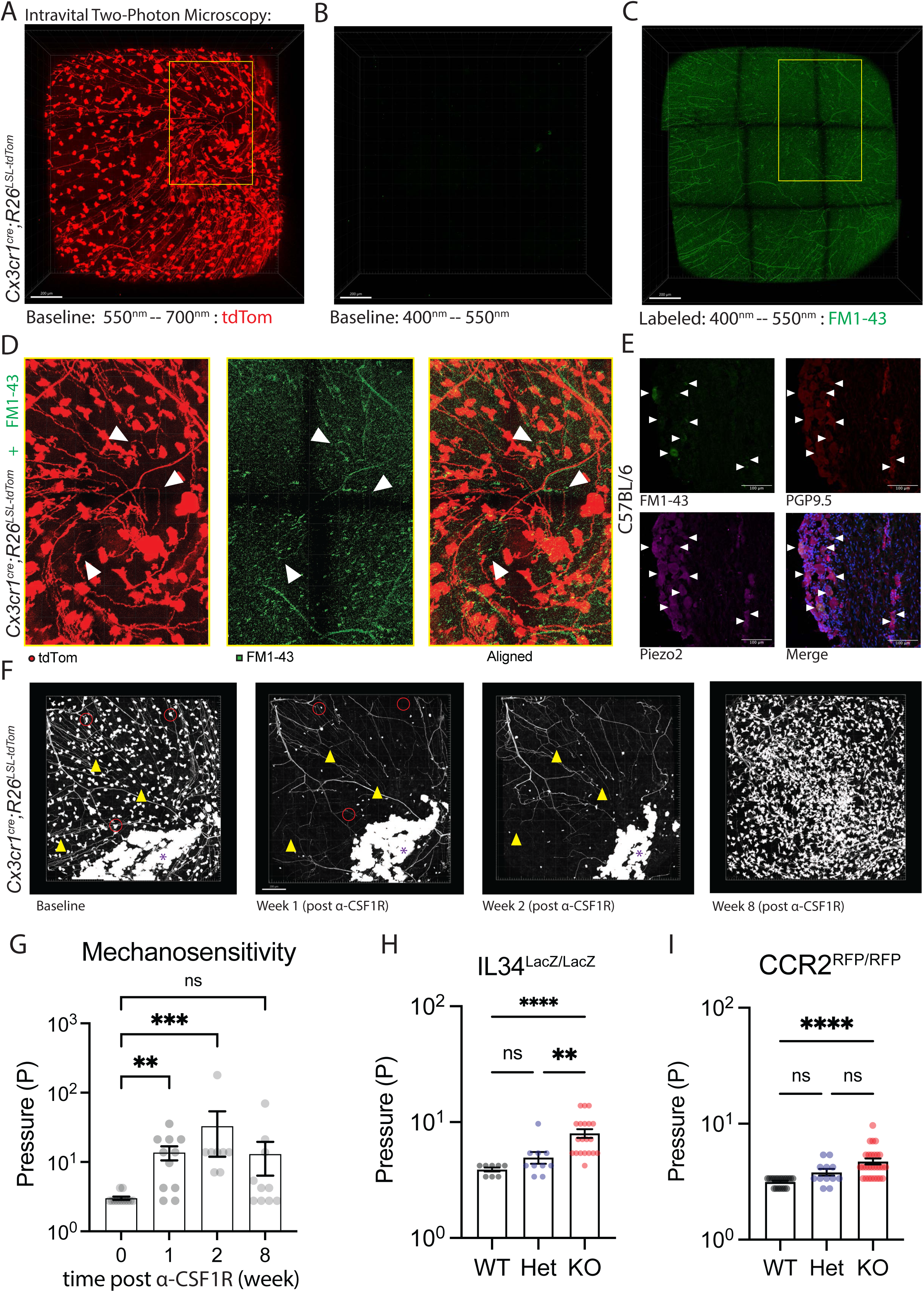
Corneal Macrophages are Critical for the Maintenance of *Piezo2*+ Corneal Nerves and Mechanosensation. (A) Baseline imaging of tdTom+ corneal nerves from *Cx3cr1^Cre^;R26^LSL-tdTom^*, visualized via intravital multiphoton microscopy (yellow box depicted in D; Scale bar = 200 µm). (B) Baseline imaging green light spectra showing no signal before labeling in corneal tissues. (C) FM1-43 labeling in the green light spectra. (D) Baseline tdTom and post FM1-43 labeling intravital images were aligned (white arrowheads depict co-labeled nerves). (E) Confocal analysis of FM1-43 labeled C57BL/6 mice shows co-labeling of Piezo2 and FM1-43 in TG somata immunolabeled for PGP9.5 and Piezo2 colocalization in FM1-43 labeled nerve population. (F) Time course intravital multiphoton imaging of the same mouse at baseline and post macrophage depletion (red circles depict depleted macrophages; yellow arrowheads highlight remodeling nerves; purple star demarcates epithelial labeling; Scale bar = 200 µm). (G) Quantification of corneal mechanosensation using Cochet-Bonnet esthesiometry following anti-CSF1R macrophage depletion (n=8-12). (H) Corneal esthesiometry in *IL34* deficient mice (n=4-10). (I) Corneal esthesiometry in *CCR2* deficient mice (n=6-14). Data shown as mean ± SEM, analyzed using one-way ANOVA and Kruskal–Wallis with multiple comparisons (∗p < 0.05; ∗∗p < 0.01; ∗∗∗p < 0.001; ∗∗∗∗p < 0.0001; ns, not significant). Data are representative from at least two independent experiments.

**Figure 6:** Schematic of the Tricellular Neuro-Immune Niche. Non-myelinating Schwann cells, which envelope the stromal nerves, maintain monocyte-derived macrophages via production of IL-34 at the epithelial basement membrane. In turn the macrophages maintain the structure and function of *Piezo2+* mechanosensitive nerves, resulting in their degeneration in the absence of corneal macrophage populations.

Having established that *Cx3cr1^Cre^* labeled nerves are functional Piezo2 mechanosensors, we sought to evaluate the functional impacts of macrophage depletion on this nerve class. Our approach employed Cochet-Bonnet esthesiometry following macrophage depletion in *Cx3cr1^Cre^; R26^LSL-tdTom^*, measuring mechanosensation. Using a nylon filament, a blink response can be elicited from mice by gentle application to the central cornea when nerves have normal function. The force applied from the filament can be increased by shortening the length of filament, indicating reduced (sometimes loss of) corneal nerve function. In addition, we captured morphological changes of tdTom+ nerves intravitally over this time course via 2P microscopy (Figure 5F). Within one week of macrophage depletion there was a significant loss of mechanosensation compared to baseline (Figure 5G). This deficit peaked around 2-4 weeks and began to re-establish near baseline levels following repletion of corneal macrophage populations by approximately the 8^th^ week (Figure 5G).

To measure the effects of macrophage depletion on C-fiber nociception, we administered eye drops with activators of TRPV1 (capsaicin), TRPM8 (Icillin or hypertonic solution) and TRPA1 (Icillin, or AITC) nociceptors, respectively. We employed an automated video capture system to quantify grimacing based on palpebral aperture in response to the eye drop stimuli (Supplemental Figure 4A). Prior to chemical behavioral, we confirmed that both control and macrophage depleted mice had equivalent palpebral apertures (Supplemental Figure 4B). Our results for TRPV1 (capsaicin [cap] eyedrops) revealed a reduced palpebral aperture following application (Supplemental Figure 4C). In contrast, we detected no significant changes following application of TRPM8/TRPA1 super-agonist (Icillin), TRPM8 (hypertonic solution), or TRPA1 (AITC) containing eyedrops (Supplemental Figures 4D-F). Our results suggest that macrophage depletion has limited effect on TRPM8 and TRPA1 function in the cornea. While we did observe that TRPV1 responses were impaired, this result may be an indirect effect given that *Trpv1* was not downregulated in *Cx3cr1^Cre^* labeled somata, and no morphological defects were observed in CGRP-GFP nerves.

Focusing on esthesiometry as the key measure of functional deficit due to macrophage depletion, we also tested genetic models of macrophage deficiency. As *Il34^Lacz/LacZ^* mice have reduced corneal macrophage numbers, we wanted to extend our analysis to determine if there was a functional effect to this deficiency. Using esthesiometry on WT, *Il34^Lacz/+^*hemizygous, *Il34^Lacz/LacZ^* deficient mice, we found significantly more pressure was required to elicit a blink response in the absence of IL-34 (Figure 5H), suggesting these mice have impaired mechanosensation. Separately, we also tested *Ccr2^RFP/RFP^*, which are *Ccr2* deficient^64^ and have a significant reduction in monocyte-derived macrophages^33^. Our results show that *Ccr2^RFP/RFP^* mice also had impaired blink reflexes as measured by esthesiometry (Figure 5I). Taken together, our results indicate that the nerves affected by macrophage deficits are Piezo2+ and mechanosensitive.

## DISCUSSION

In this study, we uncovered a previously unrecognized role for monocyte-derived macrophages in maintaining Piezo2+ mechanosensory nerve function in the cornea rendered through a site-specific and highly specialized tripartite cellular niche^60^. Our findings reveal that corneal macrophages are maintained in part by Schwann cell-derived IL-34 and specifically support Piezo2-enriched mechanosensory nerve endings. This work establishes that touch sensation in the cornea requires the orchestrated interaction of immune cells, glia, and nerves, extending our knowledge beyond the traditional neuron-centric view of mechanosensation.

To understand corneal macrophage ontogeny, we performed lineage tracing studies that highlighted the importance of monocyte-derived macrophages in populating the adult corneal stroma^29,32,33,65,29,66,67,68^. While the cornea proper is avascular, the far peripheral region (i.e., limbus) is rich in blood and lymphatic vessels, providing tissue access for circulating monocyte entry and egress. The steady replenishment of macrophages through definitive hematopoiesis aligns with the unique immunological requirements of the cornea, a tissue at a critical environmental interface. This process may allow for enhanced adaptability in response to environmental challenges ^69,70,71,72,73,74^ while maintaining precise control over nerve function.

A key discovery in our study is the identification of a specialized tripartite niche, where macrophages interact with both nerves and Schwann cells at nerve penetration sites. This anatomical arrangement appears strategically positioned to support nerve endings, as they shed their Schwann cell ensheathment and subsequently enter the corneal epithelium^75^. ScRNA-seq enrichment of nerve and glial maintenance pathways in corneal macrophages suggest that this physical co-localization has functional significance. Intriguingly, while macrophages proved critical for the maintenance of select nerves, they were dispensable for Schwann cell morphology, indicating a directed, rather than reciprocal, relationship in this three-way cellular interaction. The dependence of corneal macrophages on Schwann cell-derived IL-34 reveals a mechanism for maintaining this sensory circuit. Our findings suggest that Schwann cells, through IL-34 production, orchestrate the placement and maintenance of macrophages precisely where they are needed to support mechanosensory nerve endings. Provision of macrophage trophic support by non-immune cells has likewise been documented in the periphery, as well as the CNS^29,53,76,77^. Relatedly, Schwann cells have been shown to provide trophic support to non-neural cells^77,78^.

Another striking finding was the selective requirement of macrophages for maintaining Piezo2-enriched mechanosensory nerve endings. Although Piezo2 is expressed by both A and C fibers, several lines of evidence suggest that the affected nerves in our study are Aδ. First, while the cornea is predominantly innervated by C-fibers, only a small number of nerves were labeled with *Cx3cr1^Cre^*, indicating a specific subset. Second, most C-fibers are CGRP positive, yet we observed minimal loss of CGRP-GFP+ nerves following macrophage depletion, suggesting that the affected nerves are not the typical C-fibers. Third, behavioral responses predominantly affected mechanosensation rather than other sensory modalities, which is more characteristic of Aδ fibers. We note a reduced capsaicin-triggered behavioral response was observed in the macrophage depletion setting. However, our spatial sequencing data showed that *Cx3cr1^Cre^*-labeled somata were enriched for *Piezo2* and *Nefh*, and had downregulated expressions of *Trpv1*, *Trpa1*, and *Tac1*, further supporting the idea that these are Aδ fibers. Moreover, applying these DEGs and a gene module onto Bhuiyan *et al.* and St. Leger *et al.* datasets further suggest that *Cx3cr1^Cre^*-labeled somata represent A-fibers. The specificity of macrophage support for Aδ fibers might be related to the difference in diameter between Aδ fibers and polymodal C-fibers, with Aδ fibers being larger^19^. We hypothesize that larger fibers may require macrophage support, either mechanically or enzymatically, to effectively penetrate the dense extracellular matrix of the epithelial basement membrane. Macrophages may be just one component of a complex neuro-myeloid network that extends beyond Aδ^12,46,69,79,80^.

Taken together, our results represent a new concept in neuro-immune communication, where macrophages maintain specific populations of sensory nerves rather than providing general trophic support. Our findings establish macrophages as essential for the maintenance of Piezo2 mechanosensory nerve function and reveal a previously unrecognized complexity in the cellular interactions required for touch sensation. This work provides new insights into the fundamental mechanisms of sensory system maintenance and neuro-immune interaction.

Several questions emerge from our findings that warrant further investigation. First, the molecular mechanisms by which macrophages specifically support Piezo2-enriched nerve endings remain to be fully elucidated. Second, it is important to explore whether similar macrophage-dependent maintenance of mechanosensory neurons exists in other tissues. Finally, the potential role of this tripartite niche in disease states, particularly those affecting corneal sensitivity or mechanical pain, represents a critical area for future research.

## SUPPLEMENTAL MATERIAL

<u>Supplemental Video 1</u> – Intravital multiphoton imaging of PLP1-eGFP in homeostasis. Corneal Schwann cells are labeled with GFP (Green). Stroma was rendered using SHG (blue). Data is representative of at least n = 3 independent experiments.

<u>Supplemental Video 2</u> – Ex vivo confocal imaging of PLP1-eGFP mouse corneas. Schwann Cells are labeled with GFP (green). Nerves were labeled with Tuj1 (red). Macrophages were labeled using Cd206 (purple). Colocalization analysis was performed using the intersection of cell volumes (white). Data is representative of at least n = 3 independent experiments.

<u>Supplemental Video 3</u> – Intravital multiphoton imaging of *Cx3cr1^Cre^;R26^LSL-tdTom^* in homeostasis. Corneal nerves and macrophages are labeled with tdTom (red). Stroma was rendered using SHG (blue). Nerves were rendered as filaments (yellow). Data is representative of at least n = 3 independent experiments.

<u>Supplemental Video 4</u> – Baseline intravital multiphoton imaging of *Cx3cr1^Cre^;R26^LSL-tdTom^*, which labels corneal nerves and macrophages with tdTom (red). Stroma was rendered using SHG (blue). Nerves were rendered as filaments (yellow). Macrophages were rendered as surfaces (white). Data is representative of at least n = 3 independent experiments.

<u>Supplemental Video 5</u> – Post-depletion intravital multiphoton imaging of *Cx3cr1^Cre^;R26^LSL-tdTom^*, which labels corneal nerves and macrophages with tdTom (red). Stroma was rendered using SHG (blue). Nerves were rendered as filaments (yellow). Macrophages were rendered as surfaces (white). Data is representative of at least n = 3 independent experiments.

<u>Supplemental Video 6</u> – Intravital multiphoton imaging of WT mouse cornea labeled via i.p. injection of FM1-43, which marks Piezo2+ nerves (red). Stroma was rendered using SHG (blue).

**Supplemental Figure 1.**
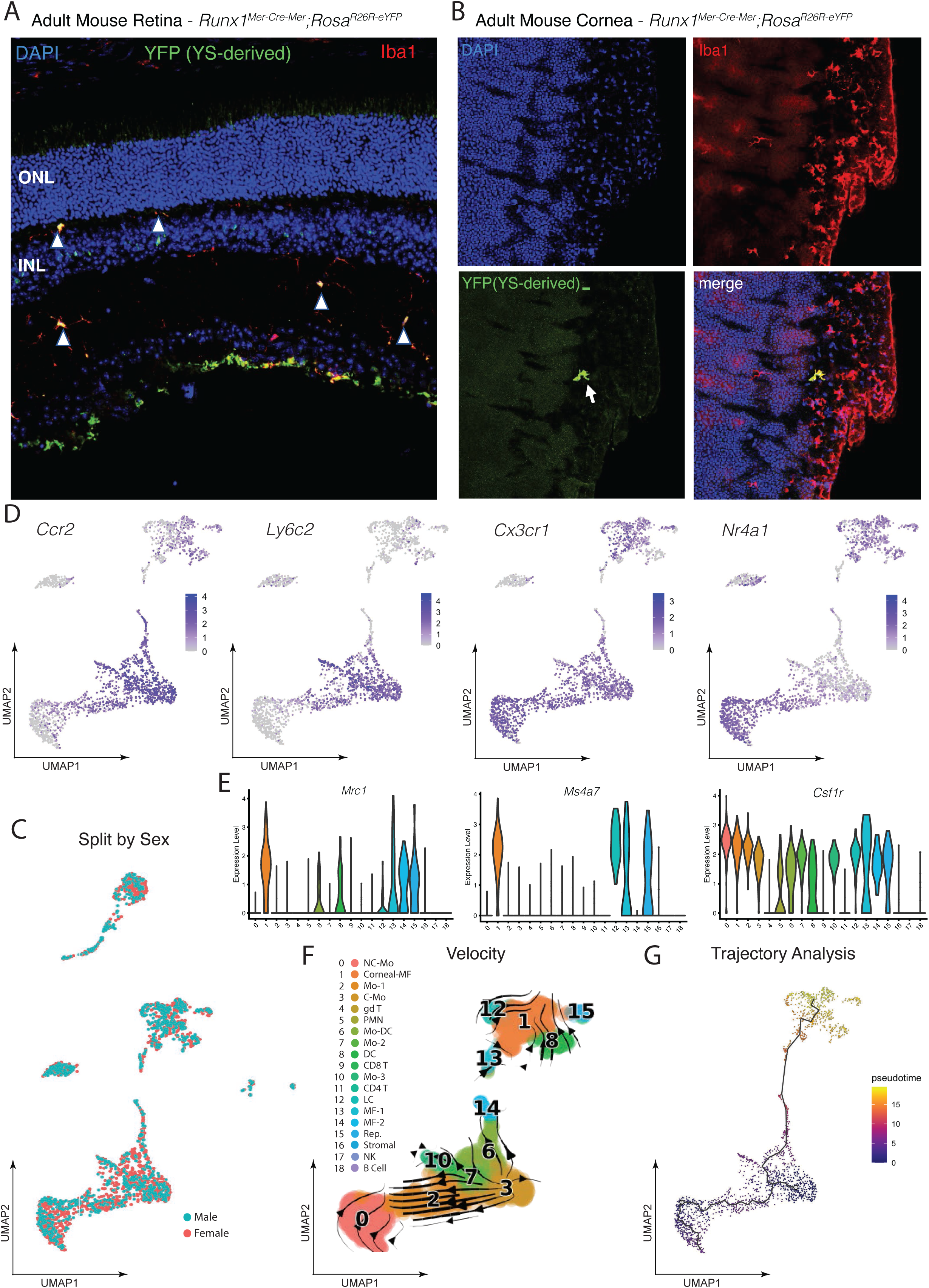
–– Lineage Tracing and Trajectory Analysis: (A) Representative images of yolk-sac-derived microglia in retina. Pregnant Runx1^Mer-Cre-Mer^; Rosa^R26R-eYFP^ mice received tamoxifen 4-OHT at E7.5. Tissues were collected from 8-week-old progeny for confocal microscopy. (B) Representative confocal microscopy images of mouse cornea from *Runx1* lineage tracing mice (C) UMAP dimensionality reduction of corneal immune cell scRNA-seq data split by sex (cyan = female; red = male). (D) Feature plots depict marker genes characteristic of circulating Classical and Non-Classical Monocytes (E) Violin plots depicting enrichment of macrophage markers. (F) RNA-velocity analysis using scVelo, showing direction of transcriptional shifts between circulating monocytes and corneal immune cell clusters. (G) Monocle3 pseudotime trajectory analysis, non-myeloid clusters were excluded from this analysis.

**Supplemental Figure 2.**
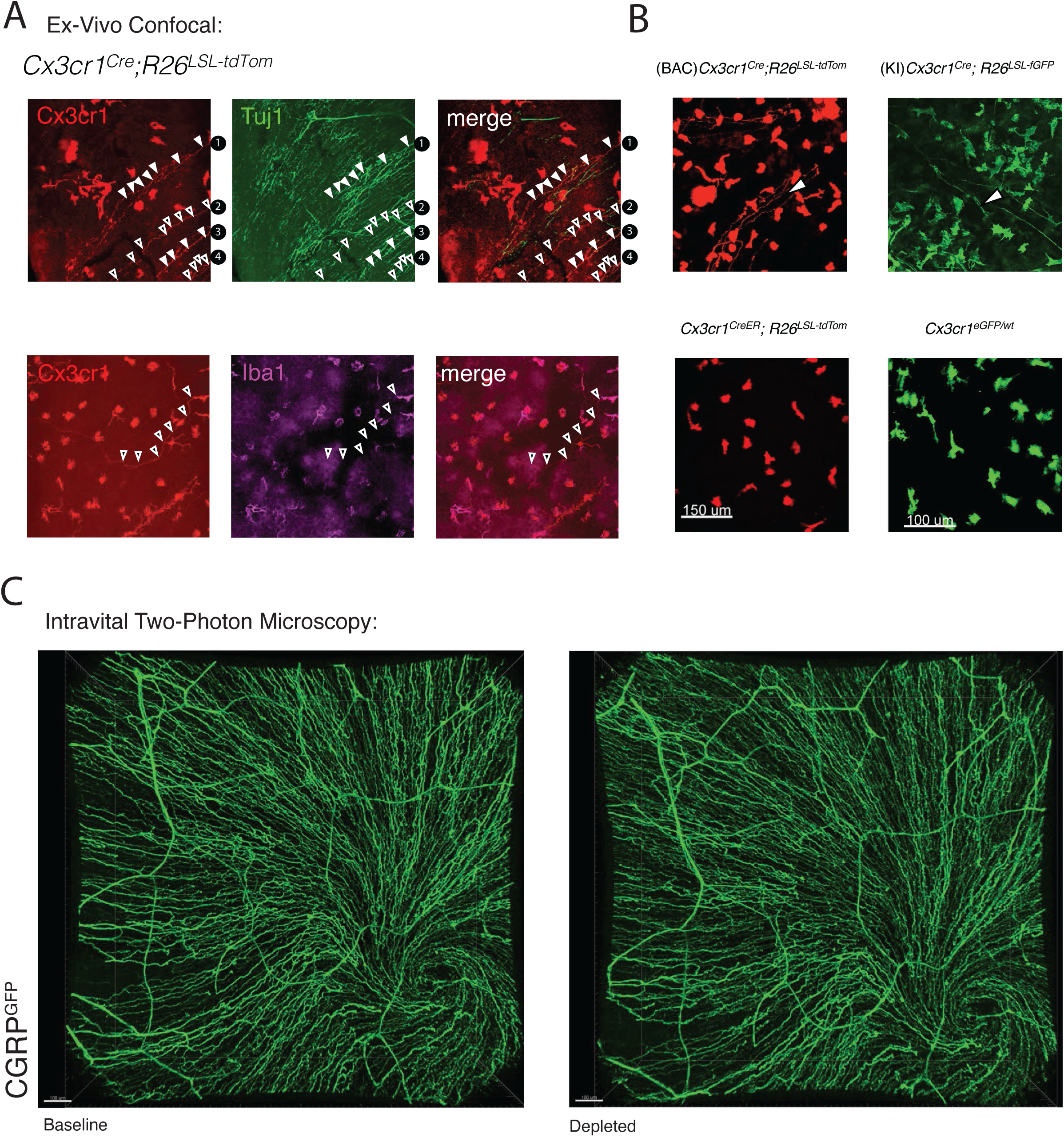
– *Cx3cr1* labels nerves in development, but not with creER: (A) Confocal microscopy of *Cx3cr1^Cre^;R26^LSL-tdTom^* corneas labeled for nerves (Tuj1, green) and macrophages (Iba1, purple). Arrowheads depict labeled nerves annotated in numerical order. (B) Confocal microscopy of *Cx3cr1^Cre^; R26^LSL-tdTom^* and knockin *Cx3cr1^Cre/+^*; *R26^LS-fGFP^* mice demonstrate dual labeling of macrophages and nerves (arrowheads). *Cx3cr1^eGFP/+^* mice and adult tamoxifen pulsed *Cx3cr1^CreERT^*^2^; *R26^LSL-tdTom^* mice lack nerve labeling (C) Intravital multiphoton imaging of CGRP-GFP mouse cornea at baseline and following macrophage depletion.

**Supplemental Figure 3.**
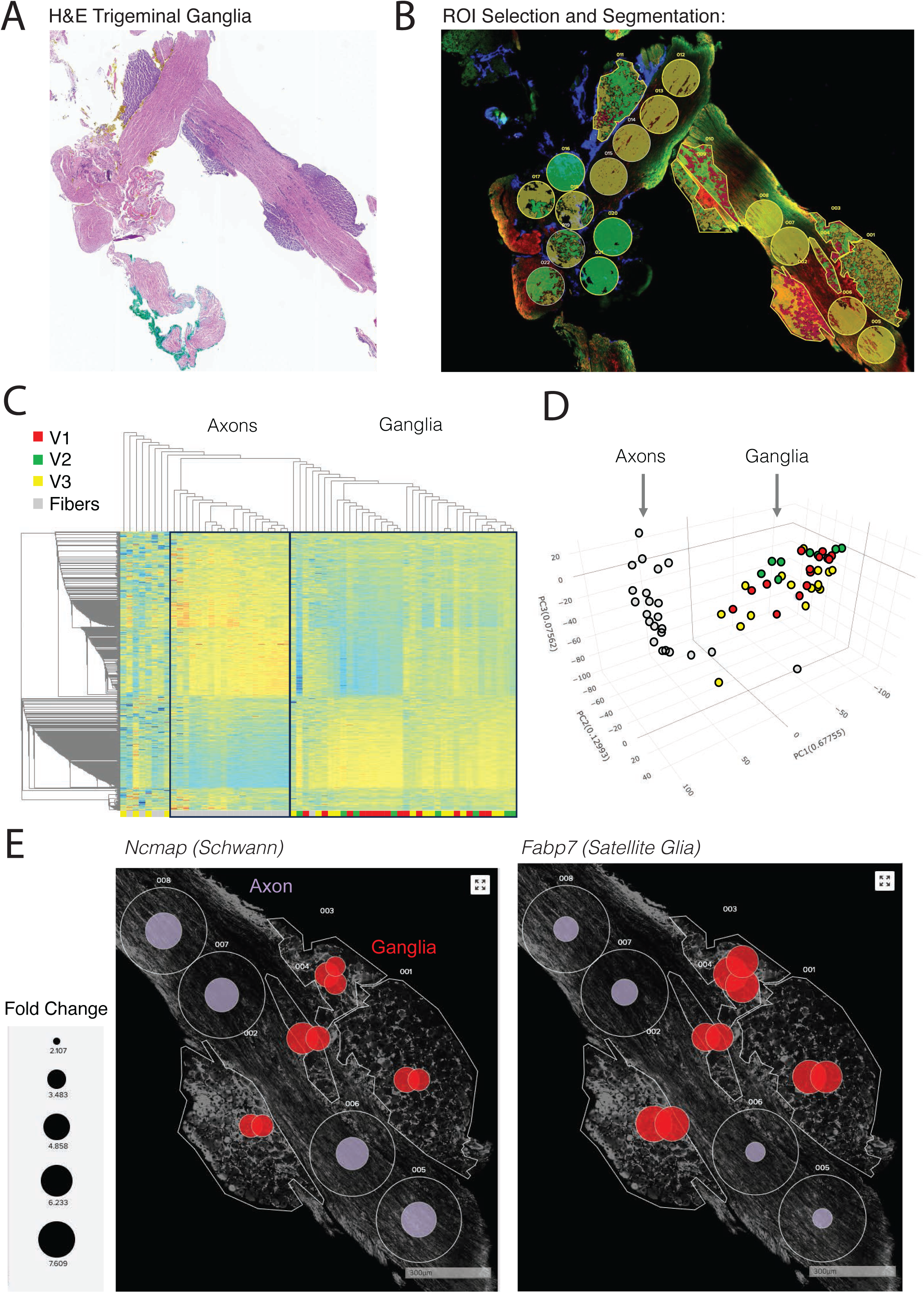
– Spatial Sequencing Analysis of Axons and Ganglia: (A) Hematoxylin and eosin (H&E) stain of sectioned trigeminal ganglia (n=2) used for spatial RNA sequencing on the GeoMx platform. (B) Representative segmentation of axons and V1, V2, and V3 ganglia regions. (C) Hierarchical clustering of axon (n=24) and ganglia (n=39) segments. (D) PCA dimensionality reduction visualizing axon vs ganglia clustering. (E) Bubble plots depicting *Ncmap* and *Fabp7* gene expression in axon and glia segments

**Supplemental Figure 4.**
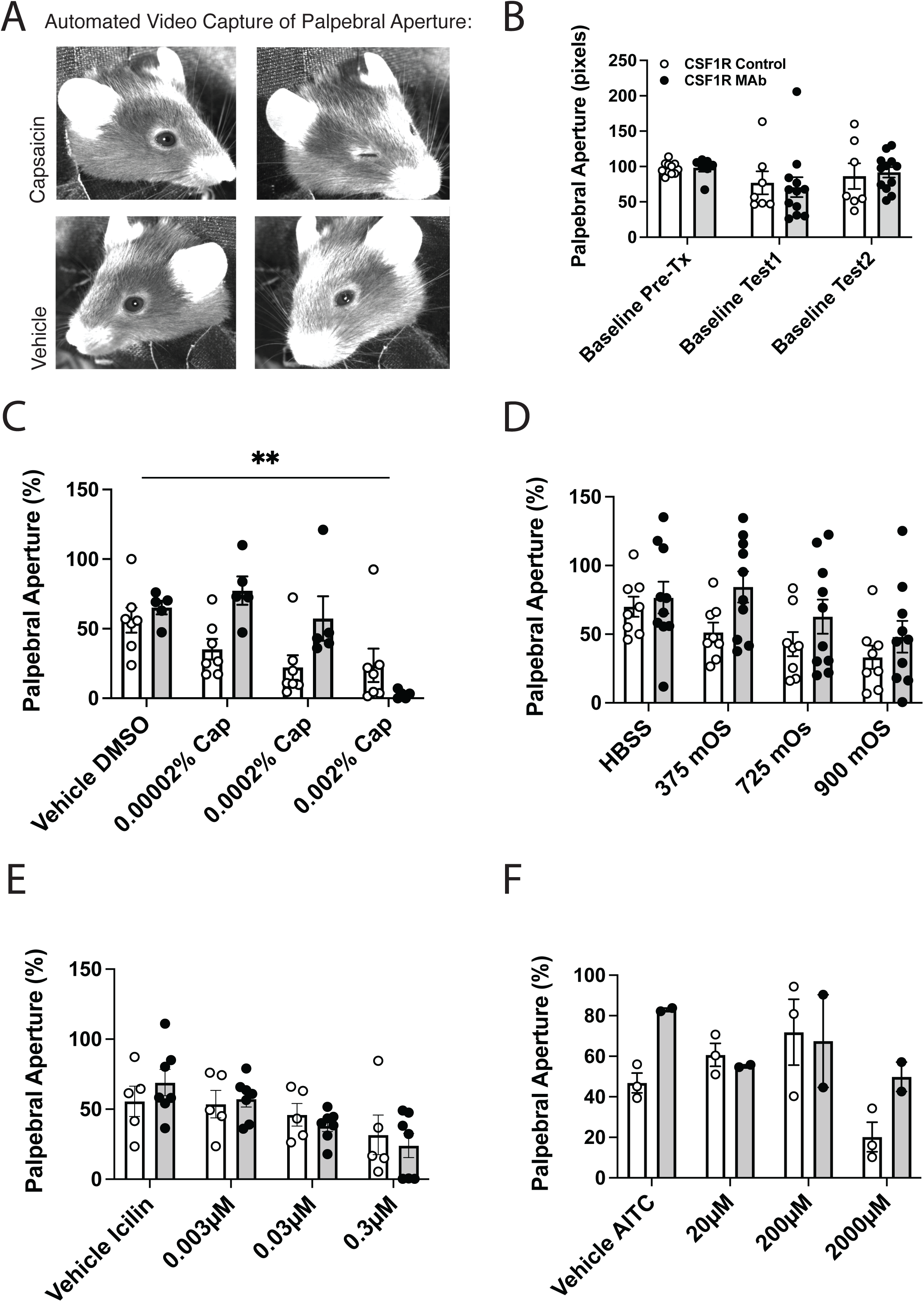
–– Quantification of Palpebral Aperture following Eyedrop Challenge: (A) Representative imaging of measurement of mouse palpebral aperture under vehicle and capsaicin, as well as baseline apertures. (B) Palpebral aperture (pixels) at baselines before treatment (macrophage depletion) and test (eyedrops). (C) Measurement of palpebral aperture following challenge with capsaicin (n=7 control, n=5 anti-CSF1R), (D) hyperosmolar (n=8 control, n=10 anti-CSF1R), (E) icillin (n=5 control, n=7 anti-CSF1R), or (F) AITC (n=3 control, n=2 anti-CSF1R).

## MATERIALS AND METHODS

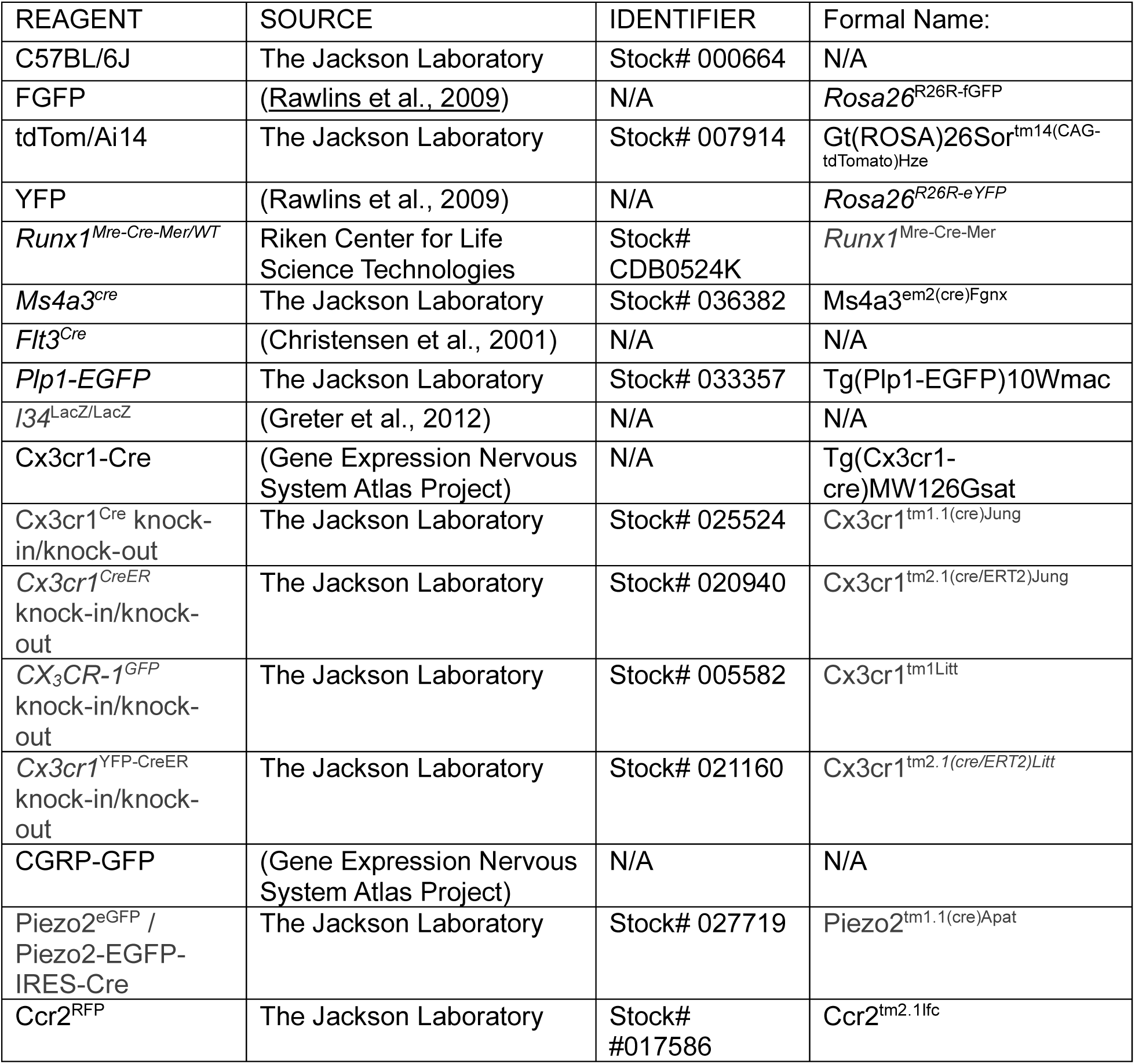

### Mice

The following mice C57BL/6J, *Cx3cr1^tm1.1(cre)Jung^*, *Cx3cr1^tm2.1(cre/ERT2)Jung^*, *Cx3cr1^tm1Litt^*, *Cx3cr1tm2.1(cre/ERT2)Litt*, as well as Tg (Plp1-EGFP)10Wmac and *Piezo2^tm1.1(cre)Apat^*, were purchased from Jackson Laboratories. Cx3cr1-Cre mice [Tg (Cx3cr1-cre) MW126Gsat also called “MW126-Cre”] were generated on a mixed 129/B6 background (Gene Expression Nervous System Atlas Project) and backcrossed for 12 generations by M.D. Gunn (Departments of Immunology and Medicine, Duke University Medical Center). *B6.Cg-Gt(ROSA)26Sor^tm^*^14^*^(CAG-tdTomato)Hze/J^* mice (referred to here as tdTom) obtained from Jackson Laboratories, were also a kind gift from M. D. Gunn. Rosa26R-CAG-fGFP mice, a kind gift from B. L. Hogan (Department of Cell Biology, Duke University Medical Center), were previously described. The *Ms4a3^Cre^* were graciously donated by Dr. Florent Ginhoux, while *Flt3^Cre^* mice were courteously supplied via Dr. Shinohara and used under MTA with Dr. T. Boehm (Max-Planck Institute of Immunobiology, Freiburg, Germany) after crossing to Rosa26 LSL-tdTomato mice. *Il34^LacZ/LacZ^*mice, generated as previously described (Greter et al., 2012), were a generous gift of Meriam Merad. CGRP-GFP were kindly provided by Dr. Perez via Alain Chédotal, and *Ccr2^RFP^* were generously provided by Dr. Edward Miao. All mice herein were bred and housed at a barrier-free and specific-pathogen-free facility at Duke University School of Medicine (Durham, NC). All procedures were approved by the <u>Institutional Animal Care and Use Committee</u> at Duke University, and the procedures were carried out in accordance with the approved guidelines. Additionally, *Runx1^Mer-Cre-Mer/WT^* and *Rosa26^R26R-eYFP/R26R-eYFP^*mice were generated and used previously as described (<u>Ginhoux et al., 2010</u>). These mice were bred and housed in the Biomedical Resource Centre, Singapore. All experiments and procedures done on these mice were approved by the Institutional Animal Care and User Committee (IACUC) of A∗STAR (Biopolis, Singapore) (Authorization No.: IACUC 151071) in accordance with the guidelines of the Agri-Food and Veterinary Authority (AVA) and National Advisory Committee for Laboratory Animal Research (NACLAR) of Singapore.

### Corneal Dissection and Whole Mount Immunohistochemistry

Mice were euthanized by CO_2_ asphyxiation before tissue harvest. Corneas were immediately dissected from mouse carcasses, or the eyes were enucleated, and placed directly in 2% paraformaldehyde (PFA) for 15 minutes at room temperature or on ice. Following 15 minutes of fixation, corneas were dissected from whole globes and subjected to an additional 45 minutes of fixation in 2% PFA. Tissues were either prepared as whole mounts or sequentially cryoprotected in 15% and 30% sucrose and then embedded in Tissue-Tek optimal cutting temperature (OCT) compound for tissue banking. Whole mounts were blocked and permeabilized with 10% serum in PBS supplemented with 0.1-1% Triton-X100 and 0.1-1% Tween-20 for either one hour at RT or overnight at 4°C. Tissues were then sequentially incubated with primary antibodies before thorough washing and incubation with appropriate secondary antibodies. Cell nuclei were counterstained with DAPI. Images were acquired with a Nikon A1 or Nikon C1/C2 confocal laser scanning microscope and analyzed using Imaris 10.1.1.

### Quantification of Whole Mount Corneal Nerves

Animals were placed under deep anesthesia with 1%-3% isoflurane, euthanized by cervical dislocation or decapitation and enucleated eyes were fixed for 20 minutes in 4% PFA in 0.1M phosphate buffer (PB) at RT. Images were captured on an Olympus Fluoview 1000 laser scanning with a 40x objective at a resolution of 1024×1024 pixels with 3µm Z-steps. A customized plugin for ImageJ was written to project the stromal, subbasal plexus and intraepithelial terminal layers separately, set thresholds mathematically for each layer and calculate the percentage of pixels containing a positive signal. Thresholds could be manually adjusted if needed. The number of Cd206-positive cells was quantitated manually in ImageJ.

### Trigeminal Ganglia Dissection and Immunohistochemistry

The trigeminal ganglia (TG) were extracted and placed in 4% PFA immediately for 1hr at 4°C and washed in PBS. For cryoprotection, the TGs were initially immersed in 15% sucrose solution until they sank at 4°C, then transferred to 30% sucrose solution and incubated overnight at 4°C to ensure thorough cryoprotection. The next day, the tissues were embedded in OCT for cryopreservation, frozen in dry ice, and stored at –80°C. Serial longitudinal slices (20 µm thick) of the TGs were obtained from dorsal to ventral using a cryostat (LEICA CM 3050 S) and picked up in Superfrost microscope slides (Thermo Fischer Scientific). The tissues were washed with cold PBS for five minutes, followed by blocking and permeabilization with 0.5% triton X (Sigma-Aldrich, St. Louise, MO) and 0.5% tween 20 (Sigma-Aldrich, St. Louise, MO) in PBS containing 5% donkey serum (blocking solution) for 1 hour. The tissues were incubated in primary antibodies overnight at 4°C. Next, tissue sections were washed in PBS four times for 10 mins and incubated with secondary antibodies for 2 hours at RT in the dark. The tissue sections were washed in PBS four times for 10 mins and stained with DAPI (Sigma-Aldrich, St. Louise, MO) for 40 minutes at RT. Followed by mounting the tissue sections in coverslips using Aqua-Poly/Mount (Polysciences) and stored at 4°C. Confocal images of the TG sections were acquired using an inverted confocal microscope (Nikon Eclipse Ti 2). Images obtained were analyzed with FIJI software (ImageJ2 2.14.0/1.54f).

The primary antibodies used were rabbit anti-Piezo2 (1:200, Novus biological, Centennial, CO) and chicken anti-PGP9.5 (1:200, Novus biological, Centennial, CO). The secondary antibodies used were donkey anti-rabbit IgG Alexa Fluor 488, donkey anti-rabbit IgG Alexa Fluor 647, donkey anti-chicken IgG Alexa Fluor 568(1:500, Jackson ImmunoResearch Laboratories, INC, West Grove, PA), and goat anti-chicken IgG Alexa Fluor 647 (1:500, Invitrogen, Thermo Fischer Scientific, Waltham, MA). The primary antibodies were diluted in the blocking solution, while the secondary antibodies and DAPI were diluted in PBS.

### Trigeminal Ganglia RNAscope

The dissected TGs were immersed in 4% PFA overnight at 4°C. For cryoprotection, the TGs were initially incubated in 15% sucrose solution until they sank at 4°C, then transferred to 30% sucrose solution and incubated overnight at 4°C. The next day, the tissues were embedded in OCT compound (Tissue-Tek) for cryopreservation. The tissues were sliced at 14 µm thickness using a cryostat (LEICA CM 3050 S) and stored at –80°C. RNAscope in situ hybridization on the fixed frozen tissue was conducted in accordance with the manufacturer’s guidelines (ACDBio, Hayward, CA). The Piezo2 probe (ACDBio, Hayward, CA) was evaluated on Piezo2cre; GFP/+ mice. For fluorescent detection, the Opal 690 reagent pack dye (1:1000, Akoya Biosciences, Marlborough, MA) was used to label the probes. All sections were mounted in coverslip using ProLong™ Diamond Antifade Mountant with DAPI (Thermo Fischer Scientific, Waltham, MA) and imaged using an inverted confocal microscope (Nikon Eclipse Ti 2). Images were analyzed with FIJI software (ImageJ2 2.14.0/1.54f).

### Intravital Multiphoton Microscopy

Mice were anesthetized via intraperitoneal injection of ketamine (100 mg/kg) and xylazine (10 mg/kg). GenTeal lubricant eye gel was applied liberally on the ocular surface to prevent desiccation. The mouse was fixed inside a light-shielded box (The Cube model CB02A, Life Imaging Services) maintained at 37 °C using a Narishige SGM-4 head holder placed on a Scientifica stage holder with breadboard platform and post on a Scientifica Motorized Movable Base Plate (MMBP). A Leica upright DM6 stand held a single objective mount for the 16x NA 0.6 HC FLUOTAR L multi-immersion objective.

Schwann Cells, nerves, and macrophages, as well as second harmonic generation (SHG) from the corneal stroma, were illuminated with a SpectraPhysics Insight X3 (680-1300 nm) or Coherent Chameleon Vision II (680-1080 nm) tunable pulsed laser using no compensation controlled by a variable beam expander and attenuated by an AOM. Emission wavelengths were discriminated via 4Tune variable dichroics and fine-tuned by continuously variable band-passes to collect adjustable ranges on HyD reflected light detectors.

Leica LAS X software (version 3.5.6.21594) was used to control parameters and collect single frames, z-stacks, and tiled images, saved as LIF files. LAS X Navigator module allowed XYZ control to search for an imaging field and collect adjacent frames for 3×3 tiled z-stacks of cornea. Resonant scanning was utilized (fixed at 8000Hz) with 4X line averaging and zoom of 1.25. Each tile was 553.57×553.57 µm (512×512 pixels, with a pixel size of 1.08 μm). Z-stack control was via objective nosepiece control in LAS X software, and the z-stack interval was calculated automatically. Z-stacks ranged from the 10µm above the corneal apex to approximately 400 µm below. The Imaris software v.9.9.1 (Bitplane AG., Belfast, UK) was used for rendering, visualization, and quantification. Volumes were quantified using Surface models. When necessary, Imaris Stitcher v.9.9.1 (Bitplane) was used to manually correct tiled image alignment.

### Tissue Harvesting and Generation of Single Cell Suspensions

Corneal tissues from adults were harvested from freshly euthanized mice. Eyes were gently enucleated, and corneas were removed under a dissection microscope. Eye tissues were further digested in Liberase TL or 1.5 mg/ml collagenase A and 0.4 mg/ml DNase I (Roche) for 1 hour at 37°C with agitation and using a Miltenyi MACS tissue dissociator. Single cell suspensions were generated by passing through 70 μm filters. Blood was collected via cardiac puncture. Cells were then treated with red blood cell lysis buffer (Sigma-Aldrich) and thoroughly washed.

### Flow Cytometry Staining and Acquisition

Single cell suspensions of cornea or lysed blood were incubated with 1% anti-FcγRIII/II clone 93 (eBioscience #14-0161-86) to block nonspecific binding. Cells were stained for 30 minutes at 4°C with a combination of fluorophore-conjugated antibodies including CD45 (BioLegend #103116), CD115 (BioLegend #135505), CD11b (BioLegend #101228), Ly6C (BioLegend #128018), and Ly6G (BioLegend #127622). Cells from blood were transferred to PBS and stained with eBioscience™ Fixable Viability Dye eFluor™ 450 (ThermoFisher #65-0863-14) for 15 minutes on ice under foil. Cells from cornea were stained with propidium iodide (Sigma-Aldrich) immediately before sorting for viability.

### Single Cell Isolation and Sorting

Corneas were isolated, with careful attention to exclude the limbus, from 25 male and 25 female WT mice at eight weeks of age. Corneas were pooled by sex and digested to generate single cell suspensions as described above. Live CD45+ corneal cells were sorted by FACS, and 2,500 to 3,000 cells of each sample were collected. An equal number of viable blood monocytes (CD45+, CD115+, CD11b+, Ly6G-) were included.

### Single Cell RNA-seq Library Preparation and Sequencing (Drop-seq)

The samples were prepared at the core facility of DMPI Molecular Genomics (Duke) using 10x Genomics Single Cell 3’ v3 chemistry for library preparation. Libraries were generated pooling cells by sex. 2,957 cells from male and 1494 cells from female mice were targeted for sequencing using a NextSeq 500 High-Output flow cell with a PE read length of 75bp by GCB Sequencing and Genomic Technologies (Duke).

### Computational Data Analysis

Sequencing data was initially processed using Cell Ranger (Version 3.0.1), samples were demultiplexed and FASTQ files were aligned to the mm10 mouse reference assembly. Feature barcode processing and unique molecular identifier (UMI) counting were performed according to standard workflows. The following criteria were applied as quality control using Seurat (version 5.10), cells that had <300 UMI counts or genes that were expressed by fewer than three cells were removed from further analysis. Cells that had >7,500 UMI counts or >10% of mitochondrial genes were also excluded. 2800 cells and 16,936 genes were used for downstream analysis. The data was scaled and normalized, and the top 2000 variable features were used to perform linear dimensionality reduction via principal component analysis (PCA).

The top 20 PCs were used to construct the Shared-Nearest Neighbor (SNN) graph and cells were clustered using the Louvain algorithm. Clusters were visualized in two dimensions via Uniform Manifold Approximation and Projection (UMAP). Clusters were annotated using Differentially Expressed Genes (DEG) and canonical markers.

### GO Pathway Enrichment Analysis

GO pathway enrichment analysis was performed using PANTHER Classification System (http://pantherdb.org/). The top 200 up-regulated genes of corneal macrophages compared with all other clusters were used.

### Publicly Available Single Cell Data

GEO accession number GSE146188 and SRA NCBI accession no. PRJNA616025 were downloaded and underwent standard processing and integration using Seurat. After filtering, the top 3,000 features were used to identify the anchors for integration, and the top 30 PCs were used to generate UMAP clustering. Clusters were labeled using marker genes identified from literature.

### Spatial Sequencing

Mice were euthanized by CO_2_ asphyxiation and perfused with PBS before tissue harvest. TGs were dissected and placed in 10% formalin for 1 hour at 4°C and stored in 70% ethanol until embedding. Total ischemic time was <15 minutes for all TG. When sufficient tissues were collected, they were embedded in paraffin. 5 µm sections were placed on glass slides for histologic examination and spatial profiling. Digital spatial whole transcriptomic profiling was performed using the GeoMx platform (NanoString/Bruker). Briefly, FFPE tissues were deparaffinized and antigen retrieval was performed using Tris-EDTA buffer (pH 9.0) at 100°C for 20 min. Tissues were post-fixed with 10% neutral buffered formalin and thoroughly washed using PBS. In-Situ hybridization with WTA probes was conducted overnight at 37°C as per manufacture instructions. Stringent washes were used to remove off-target probes. Tissues were blocked with buffer W for 30 minutes at RT, and stained for morphology markers with fluorescence-conjugated antibodies and nuclear dye for one hour at RT. Slides were loaded onto DSP for ROI collection or stored in 2x SSC at 4°C for up to two weeks. ROI were selected targeting axons and ganglia, including V1, V2, and V3 regions. ROI were segmented by cell type using nuclear stains and Tuj1 expression, as well as tdTom positive vs negative nerve somas. The GeoMx machine cleaves the unique molecular barcodes from designated ROIs and segments and collects them via microcapillary aspiration into unique wells of a 96-well plate.

Library preparation was performed per manufacturer’s instructions and sequenced using an Illumina NovaSeq 6000 or NovaSeq X. Data was analyzed according to standardized protocols using the GeoMx DSP.

### FM Styryl Dye Labeling

Unless otherwise stated, mice were injected intraperitoneally with FM 1-43FX (Invitrogen: F35355) or FM 4-64FX (Invitrogen: F34653), at a dose of 1.12 mg/kg of body weight from stocks made up in PBS, as previously described by Villarino *et al.*^26^ For immunohistochemistry analysis, the mice injected with FM 1-43FX one week before the TGs were dissected.

### Macrophage Depletion

Mice were injected intraperitoneally with 2 mg anti-CSF1R (BioXcell Clone AFS98, Cat# BE0213) in two doses at 48-hour intervals to deplete macrophage populations, as previously described by Hashimoto *et al.*^81^ Depletion was confirmed by intravital microscopy or ex-vivo confocal whole mount analysis.

### Cochet-Bonnet Esthesiometry

Corneal mechanosensation was measured with a Cochet-Bonnet esthesiometer (CBE, Western Ophthalmics, Lynwood, WA, USA). The filament was first applied in the central cornea at 60 mm (maximum length). The filament length was decreased by 5-mm increments and re-applied to the central cornea until reflex blinking was elicited.

### Tamoxifen Treatment

Tamoxifen (Sigma-Aldrich) was dissolved in corn oil to a stock concentration of 20 mg/ml. 75 mg/kg of tamoxifen was intraperitoneally injected twice with one day in between injections. Unless otherwise stated, mice were 6 to 8 weeks of age when given tamoxifen. For the study of yolk sac derived corneal macrophages, tamoxifen and progesterone were dissolved in corn oil and to a final stock concentration of 7.5 mg/ml and 3.75 mg/ml progesterone. A single dose of 60 mg/kg tamoxifen and 30 mg/kg of progesterone were injected intraperitoneally into pregnant mice at 7.5 days post-conception.

### Palpebral Aperture

Before each experiment, mice were acclimated in a dimly lit room (less than 50 lux) for at least 10 min. A customized cabinet equipped with a minimal restraint system was used to visually isolate the left and right eyes of unanesthetized mice for optimal image capture using an infrared light source and two 30 Hz cameras synchronized with custom software in MATLAB. Timestamps before eyedrop application (0-12 seconds) were classified as baseline, following the challenge, the animal was allowed to react by blinking for 10 seconds, before 60 second videos were recorded to measure PA. A rest period was allowed between trials to ensure the palpebral aperture returned to baseline. Palpebral Aperture (PA) video data was analyzed using publicly available pupillary analysis software (https://www.pupillometry.it/). Based on timestamp order, events were normalized to baseline, and those noted to have a difference in PA of 500 px^2^ within a second were labeled as artifacts and excluded from analysis.

### Pharmacological Agents/Eyedrops

Isotonic (290 mOsm/L) normal saline (Hospira NDC 0409-488-02, 0.9% Sodium Chloride Injection, USP) was prepared. Sucrose (Fisher S5-500) was added to increase tonicity to yield concentrations of 375, 725, and 900 mOsm/L. Serial dilutions of capsaicin (Sigma-Aldrich M2028-1G) solutions at 0.0002, 0.002, and 0.02% were prepared in vehicle (2% DMSO in Hanks Buffered Saline Solution). Icilin (TOCRIS 1531) in DMSO stock solution was prepared at a concentration of 8.03 x 10^-2^ M in DMSO. The solution was moderately vortexed and placed on a rotator for 1 hour. Icilin in vehicle DMSO: PBS (0.1 mg/mL) was prepared. Dilutions of icilin in PBS (pH 7.4) at 0.32 x 10^-6^, 0.64 x 10^-6^, 0.86 x 10^-6^, 1.28 x 10^-6^, and 3.21 x 10^-6^ M were prepared.

### Statistical Analysis

Graphs of single cell RNA-seq data were generated using R studio, and all other statistical graphs were generated using GraphPad Prism 7 software.

## Data Availability

The single-cell RNA and spatial sequencing data will be deposited in GEO.

## Acknowledgements

This study was funded by, NIH/NEI R01EY021798, NIH/NEI R01EY030906, NIH/NEI R01EY030864, Duke NIH Center Core Grant P30EY005722, UCLA JSEI Core Grant P30EY000331, NEI U01 Grant U01EY034687, Duke Research to Prevent Blindness (RPB) Unrestricted Grant, and RPB International Research Collaborator’s Award. We would also like to acknowledge the assistance of the Duke LMCF, Duke Cancer Institute, Duke Sequencing Core, and the Duke BioRepository and Precision Pathology Center, and Dr. Yuekan Jiao at the UCLA JSEI Microscopy and Imaging core for creating the ImageJ plugin.

## Contact for Reagent and Resource Sharing

Further information and requests for resources and reagents should be directed to and will be fulfilled by the Lead Contact, Daniel R. Saban.

